# Prophages as a source of antimicrobial resistance genes in the human microbiome

**DOI:** 10.1101/2025.03.19.644263

**Authors:** Laura K Inglis, Susanna R Grigson, Michael J Roach, Robert A Edwards

## Abstract

Prophages—viruses that integrate into bacterial genomes—are ubiquitous in the microbial realm. Prophages contribute significantly to horizontal gene transfer, including the potential spread of antimicrobial resistance (AMR) genes, because they can collect host genes.

Understanding their role in the human microbiome is essential for fully understanding AMR dynamics and possible clinical implications.

We analysed almost 15,000 bacterial genomes for prophages and AMR genes. The bacteria were isolated from diverse human body sites and geographical regions, and their genomes were retrieved from GenBank.

AMR genes were detected in 6.6% of bacterial genomes, with a higher prevalence in people with symptomatic diseases. We found a wide variety of AMR genes combating multiple drug classes. We discovered AMR genes previously associated with plasmids, such as *blaOXA-23* in *Acinetobacter baumannii* prophages or genes found in prophages in species they had not been previously described in, such *as mefA-msrD* in *Gardnerella* prophages, suggesting prophage-mediated gene transfer of AMR genes. Prophages encoding AMR genes were found at varying frequencies across body sites and geographical regions, with Asia showing the highest diversity of AMR genes.

**Importance:** Antimicrobial resistance (AMR) is a growing threat to public health, and understanding how resistance genes spread between bacteria is essential for controlling their dissemination. Bacteriophages, viruses that infect bacteria, have been recognised as potential vehicles for transferring these resistance genes, but their role in the human microbiome remains poorly understood. We examined nearly 15,000 bacterial genomes from various human body sites and regions worldwide to investigate how often prophages carry AMR genes in the human microbiome. Although AMR genes were uncommon in prophages, we identified diverse resistance genes across multiple bacterial species and drug classes, including some typically associated with plasmids. These findings reveal that prophages may contribute to the spread of resistance genes, highlighting an overlooked mechanism in the dynamics of AMR transmission. Ongoing monitoring of prophages is critical to fully understanding the pathways through which resistance genes move within microbial communities and impact human health.

## Introduction

The virome is an essential microbiome component mainly consisting of bacteriophages (hereafter referred to as phages), which are viruses infecting bacteria. Phages have two main lifecycles: the lytic lifecycle, where virulent phages hijack the host’s replication systems to create more virions, and the lysogenic life cycle, where temperate phages integrate into the bacterial host’s genome. Most temperate phages integrate into and are replicated as part of the host’s chromosome. However, lysogenic phages can also replicate extrachromosonally as phage plasmids, having characteristics of both plasmids and phages (Pfeifer & Rocha 2024; Pfeifer et al. 2022). Alternatively, they may become satellite phages that don’t encode all the necessary structural proteins and instead ‘borrow’ any required genes from other phages infecting the same host (Cohen 1983; Cohen et al. 1996). In addition, filamentous phages replicate and exit without lysing their hosts (Dehò et al. 2006; deCarvalho et al. 2023).

Phages facilitate the evolution of their hosts by mediating the transfer of other mobile genetic elements such as transposons, plasmids, and genetic islands(Xia & Wolz 2014; Christie & Dokland 2012; Touchon et al. 2017). They can carry genes that probably originated in bacteria, known as auxiliary metabolic genes, and can provide these genes to their hosts in a process called lysogenic conversion. Errors in replication and phage excision can also move genes from the bacterial genome into the phage genome and vice versa, allowing phages to collect more genes to transfer to their next hosts (Touchon et al. 2017). Horizontal gene transfer connects the microbial world in a sort of “common gene pool” (Davis & Olsen 2010). The ability of phages to confer these auxiliary metabolic genes to their bacterial hosts and facilitate the transfer of other genetic elements that new bacteria can then take up allows genes to slowly hop around the microbiome, spreading both valuable and benign genes between species (Hendrix et al. 1999).

The term auxiliary metabolic genes is somewhat of a misnomer, though, as it is well known that prophages can contain genes other than metabolic genes, virulence genes being a well-known example. Several pathogens, such as *Vibrio cholerae* (Li et al. 2003), *Corynebacterium diphtheriae* (Muthuirulandi Sethuvel et al. 2019), and *Escherichia coli* (Rodríguez-Rubio et al. 2021), get many of their toxin-production genes from their prophages. There is also evidence to suggest that prophages regularly carry antimicrobial resistance genes.

Antimicrobial resistance (AMR) is a growing public health issue. Since the discovery of penicillin in 1928, antibiotics have become indispensable for treating life-threatening infections (Hutchings et al. 2019). However, the short generation time of bacteria and the widespread use of antibiotics have led to a rapid arms race, causing the proliferation of AMR genes across various pathogens (Schmieder & Edwards 2012). Understanding how AMR genes are spread has been thoroughly researched since people became aware of the threat antibiotic resistance poses.

Since it is known that prophages can transfer bacterial genes between species, it is feasible that antimicrobial resistance (AMR) genes are transferred as well. AMR genes are common mobile genetic elements, but it was thought that the transduction events required to transfer AMR genes to and from phages were so uncommon as to be near impossible (Torres-Barceló 2018). It was not until recently that AMR genes were regularly being identified in phage genomes (Kondo et al. 2021; Pfeifer et al. 2022; Moon et al. 2020; Brown-Jaque et al. 2018; Colomer-Lluch et al. 2011). However, later studies suggest that overzealous interpretations of bioinformatics results and using lower identity thresholds may have resulted in overestimating the abundance of AMR genes in phage genomes ((Enault et al. 2017). Here, we investigate the abundance of AMR genes in the prophage regions of almost 15,000 bacteria isolated from humans. With this, we aim to determine how common phage-encode AMR genes are across the human microbiome, and in doing so, we identified phage-transferred AMR genes that were previously only mobilised by plasmids.

## Materials and Methods

### Genome selection and filtering

We retrieved 949,935 bacterial genomes from GenBank on June 1, 2022. Genomes classified as metagenome-assembled or containing more than 50 contigs were excluded to ensure high-quality assemblies. This filtering set ensured an average fragment length exceeding 60 kbp, which is longer than the upper limit for many prophages (McKerral et al. (2023)). Duplicate accessions were removed, and the genomes were curated by isolation source and isolation region based on metadata from the Pathosystems Resource Integration Center (PATRIC). The curated accessions were then filtered to remove any genomes not isolated from humans. After filtering 20,537 unique genome accessions remained, with 14,987 genomes that could be associated with one of 31 different body site categories and 5,341 genomes that could be categorised by the human host health status.

### Prophage identification

We previously identified over 5 million high-quality prophages in these genomes using PhiSpy (Akhter et al. 2012; McKerral et al. 2023), one of the most accurate prophage prediction tools, with the lowest runtime currently available (Roach et al. 2022), described in Inglis et al. (2024).

### AMR gene detection

AMR genes were identified using AMRFinder+ (version 3.10.23, database version 2021-12-21.1) (Feldgarden et al. 2021). Both nucleotide and amino acid sequences were analysed to maximise detection accuracy. Stringent cutoffs for sequence identity (≥99%) and coverage (≥99%) were applied to minimise false positives. AMR genes detected within prophage regions were further validated by comparison against the Comprehensive Antibiotic Resistance Database (CARD) and corroborated through a literature search to avoid housekeeping genes.

### Data analysis

The prevalence of AMR genes within prophage regions was compared across body sites, geographical areas, and host health status based on the PARTIC metadata to identify potential associations. Statistical analyses, including Kruskal-Wallis tests, were performed using SPSS to assess the significance of differences between AMR gene abundance across the aforementioned categories.

## Results & Discussion

### Virulence genes in prophages

Virulence genes were the most common auxiliary genes found in our prophages, with 1,683 (14.4%) of the genomes carrying a virulence gene in their prophage regions. 23.3% of nasal samples, 34.5% of skin samples, and 36.9% of urinary tract samples contained prophages with virulence genes. Most of the bacteria in these samples were *Staphylococcus aureu*s (65.3%) and *Escherichia coli* (33.7%). The most common genes were the increased serum survival protein *Iss*, nearly identical to the prophage-encoded protein *Bor*. The *Iss* protein is derived from the *Bor* protein, which is known to occur in *E. coli*’s phage λ (Johnson et al. 2008). Three of the most common genes are a staphylococcal enterotoxin, a phage-encoded staphylokinase (Bokarewa et al. 2006), and the complement inhibitor SCIN-A. The complete results list can be found in Supplementary Data 1.

### Stress genes in prophages

As defined by AMRfinder+, stress genes were the least commonly detected among our results, with only 283 genomes (2.4%) containing prophage regions that carry these genes. Stress genes were grouped into heat, metal, acid, and biocide resistance genes, including efflux pumps, transport proteins, repressors, regulators, and reductases.

Stress genes were found in the predicted prophage regions of 14/31 body sites. Stool and urine samples had the most genomes with stress genes in their prophage encoded regions, with 26.5% of bacteria whose prophages encode stress genes isolated from urine samples and 27.6% from stool. However, there were about half as many urine samples (n=1275) as stool samples (n=2967) in our dataset. While the bacterial genomes from all the body sites with more than 1000 bacterial isolates contained stress genes in their prophages, some body sites with few bacterial isolates, such as liver (n=9) and bile (n=27), also contained stress genes. The complete results list can be found in Supplementary Data 1.

93.6% of the stress genes in our prophage regions were from genomes from bacteria taken from symptomatic hosts, with only 8 genomes from bacteria from healthy or asymptomatic hosts containing stress genes. Most prophages containing stress genes were found in *E. coli*, with almost 20% of *E. coli* genomes containing them. More common bacteria in our dataset, such as *S. aureus* and *Klebsiella pneumoniae*, had comparatively fewer prophages, 0.6% and 2.6% of the respective species containing stress genes in prophage regions.

### AMR genes in prophages

The notion that phages can harbor and transmit antimicrobial resistance genes has long been debated. Originally, it was thought that phages rarely encoded AMR genes, but then, with the expansion of viromics, studies began to suggest that phages might often carry AMR genes (Modi et al. 2013). In 2017 Enault et. al. suggested that the thresholds used in these metagenomic analyses were not strict enough, showed that some of these ARGs predicted by bioinformatic methods did not confer resistance when transferred into bacteria, and that there are only two phages in the publicly available phage genomes in RefSeq that contain ARGs (Enault et al. 2017). Although they specifically analysed lytic phages, they briefly discussed how prophages are quite different, with a higher abundance of AMR genes than lytic phages (Enault et al. 2017), and that many prophages with known AMR genes have been shown to be incapable of lysis anymore.

To ensure that the AMR genes detected in this study were likely to confer a resistance phenotype, a literature search was conducted to find evidence of all the AMR genes found in predicted prophage regions (see Supplementary Data 1). We found that two genes, *fosB2* and *tet(X1)*, from three genomes were truncated gene variants that no longer produced resistance (Yang et al. 2004; Wisdom et al. 1992). Gene *ant(6)-Ia*, with 19 occurrences in our prophages, which may have unnamed variants of the gene that do not confer resistance (Rodrigues Souza et al. 2020). Gene *crpP*, with four occurrences in our prophages, did not confer the resistance to ciprofloxacin it was originally thought to but may still provide low-level fluoroquinolone resistance (Zubyk & Wright 2021).

The most common resistance genes among our prophages were the aminoglycoside nucleotidyltransferase *ANT(9)-Ia*, which is known to mediate resistance to spectinomycin (Sheng et al. 2023), the *ermB* and *tetM* genes. Although these genes are often found together (Roberts et al. 1999), we did not frequently find them together in our dataset.

### Bodysite and AMR

Different areas of the body have different microbiomes and, consequently, different viromes. We split our bacteria by the area of the human body where they were isolated to determine whether the proportion of prophages with AMR genes varied across the different body sites.

We found AMR genes in predicted prophage regions in bacteria isolated from 21 out of the 31 different areas of the body. Out of the body sites that didn’t contain AMR genes, only one site, the stomach, had more than 100 genomes in the dataset. We previously showed that the stomach largely lacked prophages (Inglis et al. 2024). In the body site with the most bacteria in our dataset, stool, over 11% of the genomes contained AMR genes on their prophages.

The number of bacterial genomes that contained prophages with AMR genes generally correlated with the total number of genomes isolated from a body site, with a few notable exceptions such as skin, tissue, and stool. Skin and tissue had many more bacterial isolates but fewer AMR genes, in our dataset than body sites with similar numbers of genomes containing AMR genes, leading to a low proportion of genomes containing prophage-encoded AMR genes in these sites. Stool, the most frequently sampled body site, had 4.7% fewer genomes with AMR-containing prophages than the second most common body site, blood, but had 134% more genomes overall. A Chi-Square test of AMR gene presence across body site showed a significant difference (p <0.001). These results suggest that while a larger sample size does allow us to find more instances of prophage regions that contain AMR genes, some areas of the body appear to have higher rates of AMR genes being found in prophages than others.

Comparing the average proportion of prophage DNA per bacteria in each body site to the amount of prophage DNA per genome (Inglis et al. 2024), we found that bacteria with AMR genes in prophage regions have, on average, higher amounts of prophage DNA in the genome than the average bacterial genome (figure 4).

**Figure 1:**
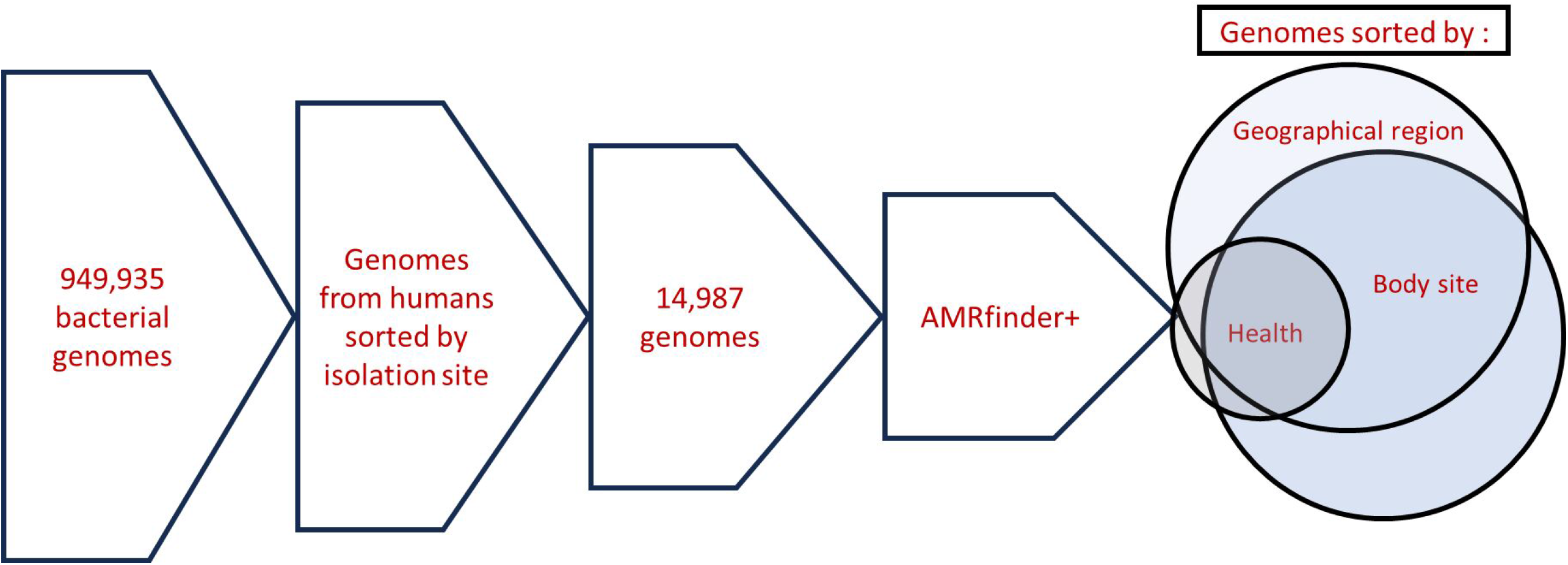
Flow chart of the methods

**Figure 2:**
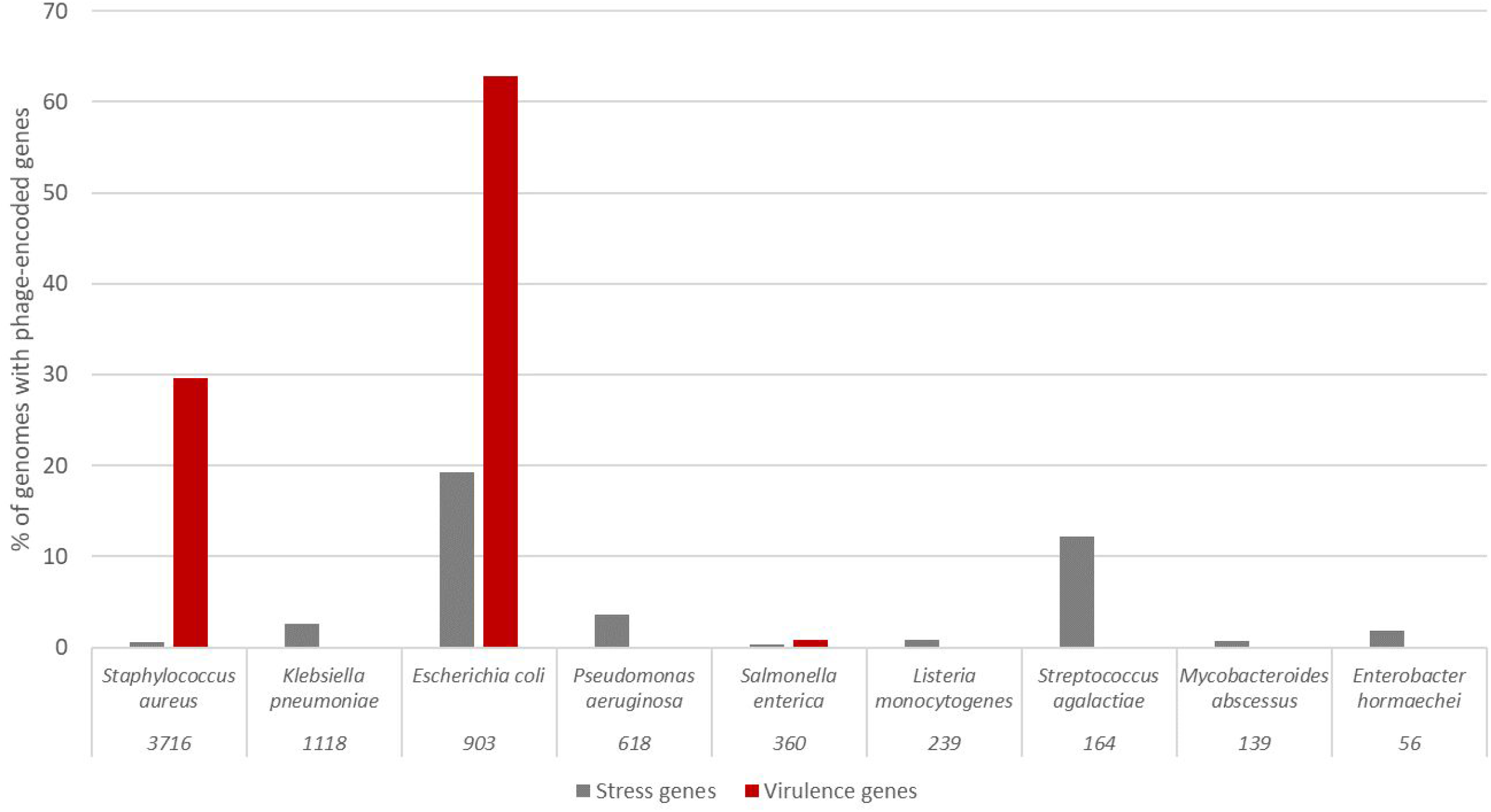
Percentage of genomes that carry either virulence or stress genes for species represented by more than 50 genomes in our dataset. The numbers beneath the species names are the number of genomes in our dataset.

**Figure 3:**
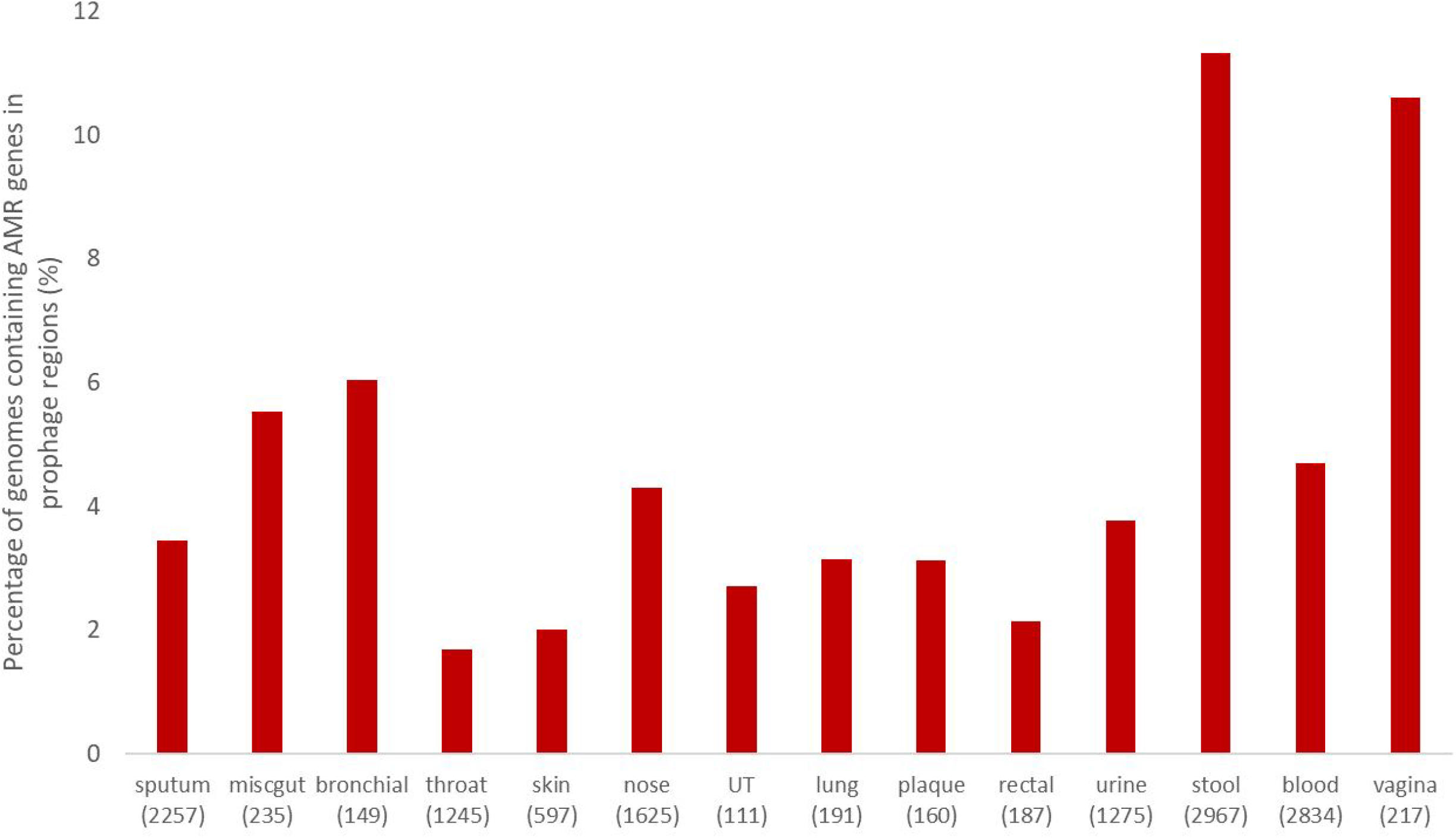
The percentage of genomes in our dataset that contained AMR genes in the predicted prophage regions. Bodysites are arranged left to right by the average amount of prophage DNA in the bacterial genome. The bracketed numbers are the number of bacterial genomes. UT and miscgut are abbreviations of urinary tract and miscellaneous gut.

**Figure 4:**
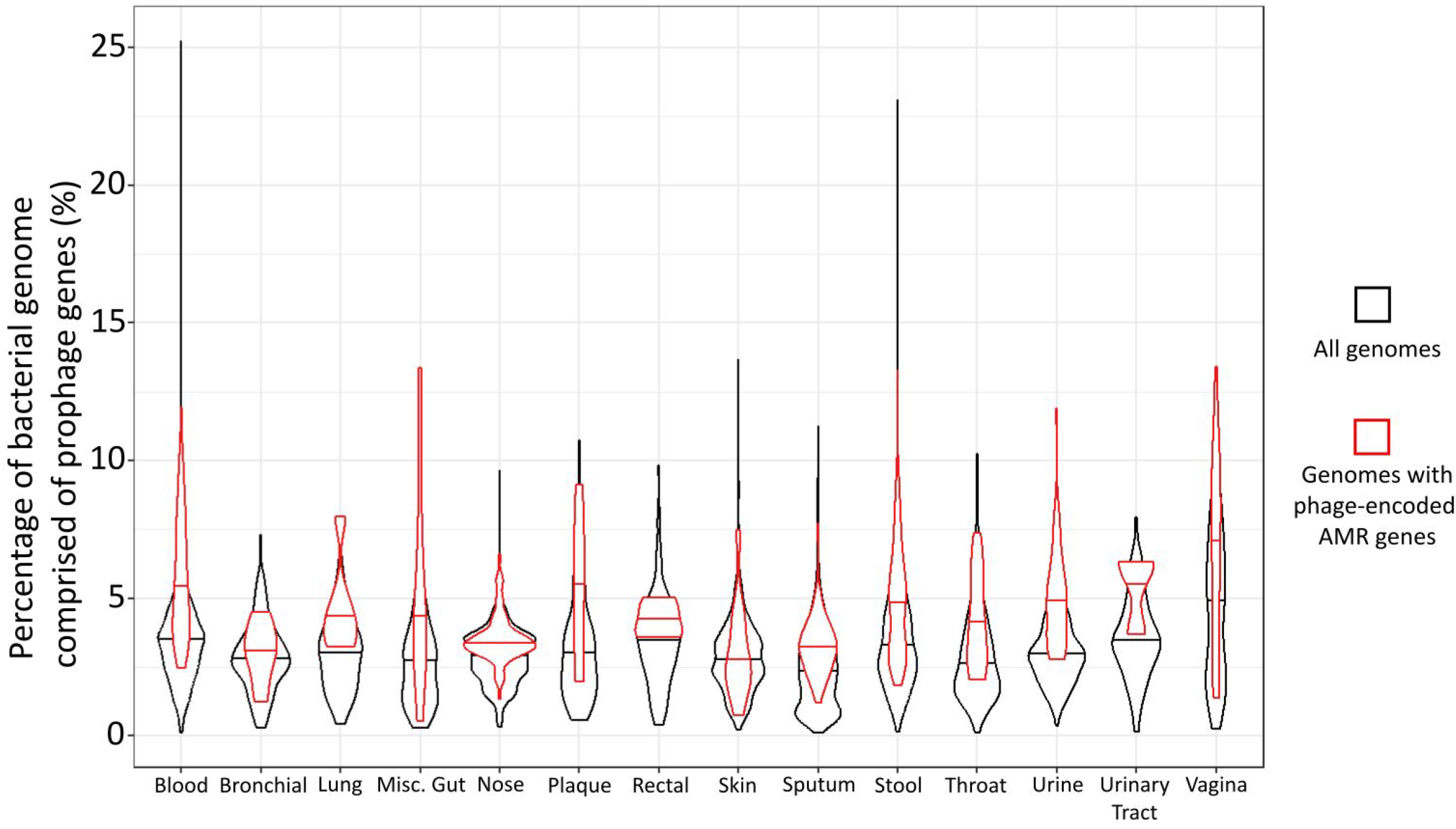
Violin plots comparing the percentage of the bacterial genome comprised of prophage DNA for all genomes in each body site (black) and genomes that contain AMR genes in their prophage regions (red). The horizontal line of the violin represents the median for each group.

There is still variation in the amount of prophage DNA the bacterial genomes contain, and it is possible for bacteria to carry multiple prophages. Therefore, we hypothesise that the bacteria carrying the phages that encode AMR genes can also still contain the non-AMR-encoding phages.

Previously, we demonstrated that stool and vaginal samples, where over 10% of prophages harbored AMR genes, ranked among the five body sites with the highest prophage DNA content (Inglis et al. 2024). While our results suggest that the likelihood of bacteria containing prophage-encoded AMR genes correlates with the amount of prophage DNA present, a Mann-Whitney U test showed there is no significant correlation (p=0.303) between the presence of prophage-encoded AMR genes and total genome size.

### Host Geography and AMR

The composition of the human microbiome and the prophages within them vary with geographical location (Suzuki & Worobey 2014; Yatsunenko et al. 2012; Gaulke & Sharpton 2018; Inglis et al. 2024), and countries have varied rates of antibiotic consumption (Klein et al. 2018).

Most geographic regions had AMR genes in the prophage regions of 3-7% of their genomes. South America had the lowest proportion of genomes with AMR genes on their prophage regions at 2.3%. Africa and Asia, with 8.4% and 9.7% of the genomes, respectively, were the most likely to have AMR genes in their prophage regions

Asia also has the most different kinds of AMR genes, with 73 unique genes across 17 classes, and North America has the second most, with 72 genes across 14 classes (figure 5). The number of unique genes mostly aligns with the sample size, with regions with more genomes submitted having a wider variety of AMR genes represented. There were two exceptions to this trend, Africa and Asia. Africa had slightly more genomes than South America, 663 and 659 genomes, respectively, but South American prophages have almost double the rate of AMR genes than those from Africa. Asia has the most unique AMR genes but the third highest number of genomes and approximately half the number of genomes as from North America. None of the genes were found in prophages in every region.

**Figure 5:**
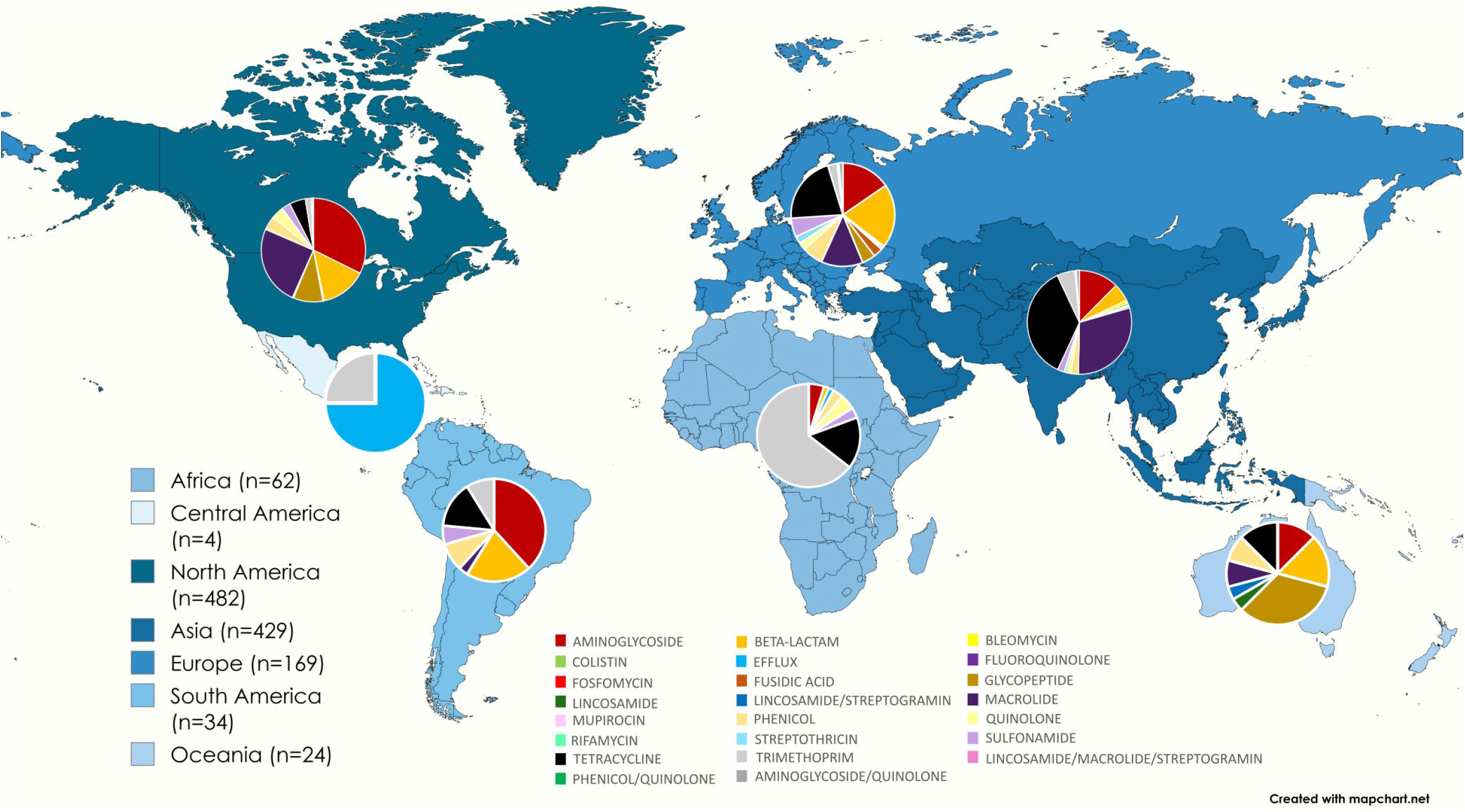
A map showing the AMR genes found in the prophage regions of genomes from each region split by drug class as defined by AMRfinder+, with n being the total number of genes found in each region.

A Kruskal-Wallis test showed some significant differences in the average number of AMR genes in prophage regions per genome. Asia, North America, and Africa were all significantly different from each other (p= <0.001-0.002). Samples from Asia were the most likely to contain AMR genes in phage regions, with 9.7% of genomes harbouring prophages that carried AMR genes. The least likely regions to find bacteria with prophage-encoded AMR genes are South America (2.3%) and Europe (2.7%).

All regions have a large variety of AMR classes, with Europe having the most classes represented (figure 5). Resistance to beta-lactams was the most common in all regions except for Africa, with aminoglycoside resistance being slightly more common, and Oceania, with efflux pumps being most common.

### Host health and AMR

There was no significant difference in the number of prophage AMR genes per genome between healthy, asymptomatic, and symptomatic samples (Kruskal-Wallis p= 0.409). However, more genomes were sampled from symptomatic humans than asymptomatic humans, and symptomatic samples were isolated from a wider variety of body sites and geographical regions. A wide variety of AMR genes was found in those samples. 6.6% of the 4,585 genomes from symptomatic hosts contained a predicted prophage with an AMR gene.

The genomes of the 159 bacterial isolates from healthy hosts contained fewer genes, 5.7% of which contained AMR genes in predicted prophage regions. The 609 genomes labeled as coming from asymptomatic individuals had a similar proportion of genomes with AMR genes (2.6%).

As bacterial genomes from healthy controls were much rarer in the GenBank database than genomes from symptomatic humans, the results we can draw from the comparisons are limited by low sample sizes. However, *Streptococcus pneumoniae* had more samples labeled as asymptomatic than symptomatic, and it showed a similar trend to other results, with bacteria isolated from asymptomatic hosts being less likely to have AMR genes in their prophage regions than bacteria isolated from symptomatic hosts. The difference is much more pronounced than most of the other species. However, bacteria from asymptomatic hosts are less likely to carry prophages with AMR genes than bacteria from symptomatic hosts. Antibiotics are given to treat illnesses and are thus more likely to be given to symptomatic humans, so there is less selective pressure for antibiotic resistance in healthy populations.

Out of the eight bacterial species that were sampled from both asymptomatic and symptomatic humans, seven species contained AMR genes in their phage regions (figure 6), while one species, *Bacteroides fragilis*, contained no AMR genes on prophages at all. Two species, *Escherichia coli* and *Staphylococcus epidermidis*, contained no AMR genes in the asymptomatic samples. One species, *Clostridioides difficile*, formerly known as *Clostridium difficile*, has a higher percentage of prophage genomes from asymptomatic hosts having AMR genes than prophage genomes from symptomatic hosts. The remaining four have a lower rate of genomes with AMR genes in prophages in asymptomatic samples than in symptomatic samples.

**Figure 6:**
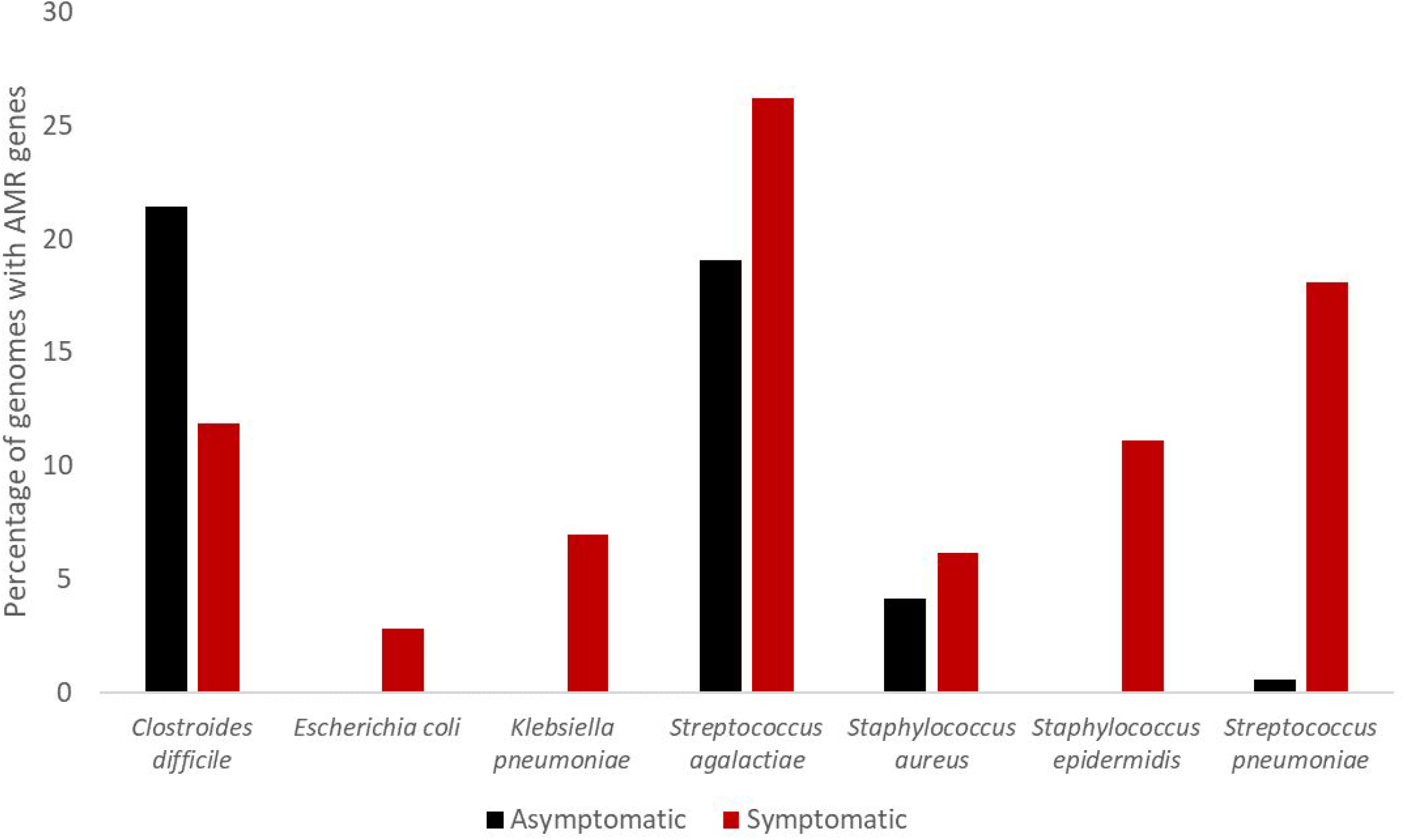
Percentage of bacterial genomes containing AMR genes in their prophage regions for seven species, split by the health of the human host (red: hosts that are symptomatic for illness, black: hosts that have no symptoms).

### Known phage-encoded AMR genes

Our dataset contained a wide variety of AMR genes. These included genes previously known to be found in prophages, such as *MefA-MsrD*, and some that were only previously known to occur in other mobile genetic elements, such as *BlaOXA23* and *TMexCD-ToprJ*.

Fosfomycin resistance has been known to be encoded by multiple lytic phage isolates of γ phage (Schuch & Fischetti 2006), and it was thought that this *FosB* gene originally came from a prophage as the wild type Wβ phage does not carry the gene (Schuch & Fischetti 2006; Gillis & Mahillon 2014). While we did not find a prophage containing the *FosB* gene, we did find prophages infecting the same species, *Bacillus anthracis*, that carried the spliced variant of the *fosB* gene, *fosB*2, even though *fosB2* cannot transform fibroblasts without the presence of a trans-activation domain as it has lost its activation domain (Wisdom et al. 1992).

The *TMexCD-ToprJ* gene clusters have been shown to confer resistance to carbapenems and tigecycline, a last-resort antibiotic for carbapenem-resistant bacteria (Zhu et al. 2023; Lv et al. 2020). This gene cluster has been identified in multiple Pseudomonas species, and we found it in *Pseudomonas aeruginosa*, both in the bacterial genomes and in a prophage region in genome GCA_008195485.1 collected in 2001.

The beta-lactamase *BlaOXA23* is a major source of carbapenem resistance for *Acinetobacter* species. Previously, *BlaOXA23* was found on conjugative plasmids (Zong et al. 2020), though few *Acinetobacter* species have conjugative plasmids, which was proposed to limit the transmission of this gene. The same penem resistance gene has also been shown to be found on transposons (Zong et al. 2020). However, we found three instances of the beta-lactamase in prophage regions of *Acinetobacter baumannii*. Phage-mediated transfer of beta-lactamases is potentially concerning as it suggests that there has been at least one instance of an AMR gene being transferred from a plasmid to a prophage. They were the only instances of AMR genes found in prophage regions of *A. baumannii*.

Phages are known to be able to transfer the macrolide efflux pump encoded by the *MefA-MsrD* gene pair to their *Streptococcus pneumoniae* hosts (Fox et al. 2021), however, we also found it in a *Gardnerella spp*. prophage.

ESKAPEE pathogens are a group of species, specifically known for their high rates of antibiotic resistance. They also face increased selection pressure due to the amount of antibiotics used to eliminate these multidrug-resistant infections. We found that while the AMR genes were very common among the ESKAPE species in our dataset, the rates of AMR genes occurring on their prophages were not above average. We hypothesise that prophages don’t face as much selection pressure as their hosts to gather resistance genes. However, there were cases where either a gene was unique to prophages or a problematic AMR gene was shown to have moved to a prophage. For example, *Ant(6)-Ia* was found in many species, both in prophage regions and in the bacterial DNA, but in *Streptococcus pyogenes*, it was only found in the prophage regions

A concern in the fight against AMR is that genes can be transferred between bacteria by horizontal transmission, spreading resistance to previously susceptible bacteria. We found 34 bacterial species, each represented more than once in our dataset, that contained specific AMR genes *only* on prophage regions. However, homologous genes could also be found in both prophage and bacterial regions in the genomes of other species, suggesting that the prophages are mobilising these genes.

While most of our common species had prophages that contained AMR genes even if they were quite rare, there were species in our dataset that did not contain any prophages with AMR genes at all, suggesting that not all phages have equal access to the common gene pool and antimicrobial resistance genes.

## Conclusion

After analysing the AMR genes in almost 15,000 bacterial genomes we found that AMR genes were relatively rare in human prophages, however, there was a wide variety of them. Different areas of the body and regions of the globe both had significantly different rates of AMR genes appearing on prophages and differing amounts of unique AMR genes occurring. Host health was only a significant factor when examining specific species. *S. pneumoniae* showed a significant difference between the number of asymptomatic and symptomatic genomes containing phage-encoded AMR genes, and three species, *E. coli, K. pneumoniae*, and *S. epidermidis*, contained no phage-encoded AMR genes in asymptomatic samples. Our results suggest that bacterial genomes with larger or multiple prophages were more likely to have a prophage that carries an AMR gene.

While AMR genes being found in prophage regions are currently rare, we identified at least one example of a gene that was known to be plasmid-bourne being found on a prophage, an instance of a gene that has moved between different prophages, of different life cycles, some genomes that only have AMR genes in their prophage regions, and genomes that appear to have acquired genes by prophages.

## Acknowledgments

We would like to acknowledge the support provided by Flinders University for HPC research resources (Flinders University 2021).

We would also like to acknowledge the use of ChatGPT to assist with grammar and layout tips.

## Supplementary Materials

Supplementary Data 1-Inglis, Laura (2025). List of unique antimicrobial resistance genes found in prophage regions. Flinders University. Dataset. https://doi.org/10.25451/flinders.28282070.v1

## Data Availability

Full AMRfinder+ results table for prophage regions is available at: Inglis, Laura (2025). AMRfinder+ results. Flinders University. Dataset. https://doi.org/10.25451/flinders.28282106.v1

## Reference list

Akhter, S., Aziz, R.K. & Edwards, R.A., 2012. PhiSpy: a novel algorithm for finding prophages in bacterial genomes that combines similarity- and composition-based strategies. Nucleic acids research, 40(16), p.e126.

Bokarewa, M.I., Jin, T. & Tarkowski, A., 2006. Staphylococcus aureus: staphylokinase. The international journal of biochemistry & cell biology, 38(4), pp.504–509.

Brown-Jaque, M. et al., 2018. Antibiotic resistance genes in phage particles isolated from human faeces and induced from clinical bacterial isolates. International journal of antimicrobial agents, 51(3), pp.434–442.

Christie, G.E. & Dokland, T., 2012. Pirates of the Caudovirales. Virology, 434(2), pp.210–221.

Cohen, G., 1983. Electron microscopy study of early lytic replication forms of bacteriophage P1 DNA. Virology, 131(1), pp.159–170.

Cohen, G. et al., 1996. The bacteriophage P1 lytic replicon: directionality of replication and cis-acting elements. Gene, 175(1-2), pp.151–155.

Colomer-Lluch, M., Jofre, J. & Muniesa, M., 2011. Antibiotic resistance genes in the bacteriophage DNA fraction of environmental samples. PloS one, 6(3), p.e17549.

Davis, J.J. & Olsen, G.J., 2010. Modal Codon Usage: Assessing the Typical Codon Usage of a Genome. Molecular biology and evolution, 27(4), pp.800–810.

deCarvalho, T. et al, 2023. Simultaneous entry as an adaptation to virulence in a novel satellite-helper system infecting Streptomyces species. The ISME journal, 17(12), pp.2381–2388.

Dehò, G., Ghisotti, D. & Others, 2006. The satellite phage P4. The bacteriophages, 2, p.391.

Enault, F. et al., 2017. Phages rarely encode antibiotic resistance genes: a cautionary tale for virome analyses. The ISME journal, 11(1), pp.237–247.

Feldgarden, M. et al., 2021. AMRFinderPlus and the Reference Gene Catalog facilitate examination of the genomic links among antimicrobial resistance, stress response, and virulence. Scientific reports, 11(1), p.12728.

Flinders University, 2021. Deep Thought (HPC). Available at: https://deepthoughtdocs.flinders.edu.au/en/latest/.

Fox, V. et al., 2021. Predicted transmembrane proteins with homology to Mef(A) are not responsible for complementing mef(A) deletion in the mef(A)-msr(D) macrolide efflux system in Streptococcus pneumoniae. BMC research notes, 14(1), p.432.

Gaulke, C.A. & Sharpton, T.J., 2018. The influence of ethnicity and geography on human gut microbiome composition. Nature medicine, 24(10), pp.1495–1496.

Gillis, A. & Mahillon, J., 2014. Phages preying on Bacillus anthracis, Bacillus cereus, and Bacillus thuringiensis: past, present and future. Viruses, 6(7), pp.2623–2672.

Hendrix, R.W. et al., 1999. Evolutionary relationships among diverse bacteriophages and prophages: all the world’s a phage. Proceedings of the National Academy of Sciences of the United States of America, 96(5), pp.2192–2197.

Hutchings, M.I., Truman, A.W. & Wilkinson, B., 2019. Antibiotics: past, present and future. Current opinion in microbiology, 51, pp.72–80.

Inglis, L.K., Roach, M.J. & Edwards, R.A., 2024. Prophages: an integral but understudied component of the human microbiome. Microbial genomics, 10(1). Available at: 10.1099/mgen.0.001166.

Johnson, T.J., Wannemuehler, Y.M. & Nolan, L.K., 2008. Evolution of the iss Gene in Escherichia coli. Applied and environmental microbiology, 74(8), pp.2360–2369.

Klein, E.Y. et al., 2018. Global increase and geographic convergence in antibiotic consumption between 2000 and 2015. Proceedings of the National Academy of Sciences of the United States of America, 115(15), pp.E3463–E3470.

Kondo, K., Kawano, M. & Sugai, M., 2021. Distribution of Antimicrobial Resistance and Virulence Genes within the Prophage-Associated Regions in Nosocomial Pathogens. mSphere, 6(4), p.e0045221.

Li, M. et al., 2003. Comparative genomic analyses of the vibrio pathogenicity island and cholera toxin prophage regions in nonepidemic serogroup strains of Vibrio cholerae. Applied and environmental microbiology, 69(3), pp.1728–1738.

Lv, L. et al., 2020. Emergence of a Plasmid-encoded resistance-nodulation-division efflux pump conferring resistance to multiple drugs, including tigecycline, in Klebsiella pneumoniae. mBio, 11(2). Available at: https://journals.asm.org/doi/abs/10.1128/mbio.02930-19.

McKerral, J.C. et al., 2023. The Promise and Pitfalls of Prophages. bioRxiv : the preprint server for biology. Available at: 10.1101/2023.04.20.537752.

Modi, S.R. et al., 2013. Antibiotic treatment expands the resistance reservoir and ecological network of the phage metagenome. Nature, 499(7457), pp.219–222.

Moon, K. et al., 2020. Freshwater viral metagenome reveals novel and functional phage-borne antibiotic resistance genes. Microbiome, 8(1), p.75.

Muthuirulandi Sethuvel, D.P. et al., 2019. Insights to the Diphtheria Toxin Encoding Prophages amongst Clinical Isolates of Corynebacterium diphtheriae from India. Indian journal of medical microbiology, 37(3), pp.423–425.

Pfeifer, E., Bonnin, R.A. & Rocha, E.P.C., 2022. Phage-plasmids spread antibiotic resistance genes through infection and lysogenic conversion. mBio, 13(5), p.e0185122.

Pfeifer, E. & Rocha, E.P.C., 2024. Phage-plasmids promote recombination and emergence of phages and plasmids. Nature communications, 15. Available at: https://www.nature.com/articles/s41467-024-45757-3.

Roach, M.J. et al., 2022. Philympics 2021: Prophage Predictions Perplex Programs. F1000Research, 10(758), p.758.

Roberts, M.C. et al., 1999. Nomenclature for macrolide and macrolide-lincosamide-streptogramin B resistance determinants. Antimicrobial agents and chemotherapy, 43(12), pp.2823–2830.

Rodrigues Souza, S.S. et al., 2020. Occurrence and associated characteristics of a mutatedant(6’)-Iagene amongEnterococcus faeciumstrains expressing phenotypic susceptibility to high levels of streptomycin. bioRxiv. Available at: https://www.biorxiv.org/content/10.1101/2020.12.28.424548.abstract.

Rodríguez-Rubio, L. et al., 2021. Bacteriophages of Shiga Toxin-Producing Escherichia coli and Their Contribution to Pathogenicity. Pathogens, 10(4). Available at: 10.3390/pathogens10040404.

Schmieder, R. & Edwards, R., 2012. Insights into antibiotic resistance through metagenomic approaches. Future microbiology, 7(1), pp.73–89.

Schuch, R. & Fischetti, V.A., 2006. Detailed genomic analysis of the Wbeta and gamma phages infecting Bacillus anthracis: implications for evolution of environmental fitness and antibiotic resistance. Journal of bacteriology, 188(8), pp.3037–3051.

Sheng, X. et al., 2023. ANT(9)-Ic, a novel chromosomally encoded aminoglycoside nucleotidyltransferase from Brucella intermedia. Microbiology spectrum, 11(3), p.e0062023.

Suzuki, T.A. & Worobey, M., 2014. Geographical variation of human gut microbial composition. Biology letters, 10(2), p.20131037.

Torres-Barceló, C., 2018. The disparate effects of bacteriophages on antibiotic-resistant bacteria. Emerging microbes & infections, 7(1), p.168.

Touchon, M., Moura de Sousa, J.A. & Rocha, E.P., 2017. Embracing the enemy: the diversification of microbial gene repertoires by phage-mediated horizontal gene transfer. Current opinion in microbiology, 38, pp.66–73.

Wisdom, R. et al., 1992. Transformation by FosB requires a trans-activation domain missing in FosB2 that can be substituted by heterologous activation domains. Genes & development, 6(4), pp.667–675.

Xia, G. & Wolz, C., 2014. Phages of Staphylococcus aureus and their impact on host evolution. Infection, genetics and evolution: journal of molecular epidemiology and evolutionary genetics in infectious diseases, 21, pp.593–601.

Yang, W. et al., 2004. TetX is a flavin-dependent monooxygenase conferring resistance to tetracycline antibiotics. The journal of biological chemistry, 279(50), pp.52346–52352.

Yatsunenko, T. et al., 2012. Human gut microbiome viewed across age and geography. Nature, 486(7402), pp.222–227.

Zhu, J. et al., 2023. Identification of TMexCD-TOprJ-producing carbapenem-resistant Gramnegative bacteria from hospital sewage. Drug resistance updates: reviews and commentaries in antimicrobial and anticancer chemotherapy, 70(100989), p.100989.

Zong, G. et al., 2020. The carbapenem resistance gene blaOXA-23 is disseminated by a conjugative plasmid containing the novel transposon Tn6681 in Acinetobacter johnsonii M19. Antimicrobial resistance and infection control, 9(1), p.182.

Zubyk, H.L. & Wright, G.D., 2021. CrpP is not a fluoroquinolone-inactivating enzyme. Antimicrobial agents and chemotherapy, 65(8), p.e0077321.

